## Supplementary Section/Summary for "Perceptual restoration fails to recover unconscious processing for smooth eye movements after occipital stroke"

**Supplementary Materials**

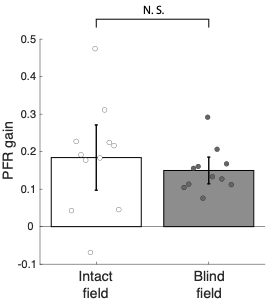
**Occipital stroke patients retain functional, smooth following eye movements at the fovea.** We found that when stimulus motion was presented foveally after the saccade on trials where the stimulus remained present, patients exhibited following movements for both intact and blind visual fields. Following was evidenced by a positive PFR gain that emerged after a visuomotor delay of about 100 ms (Figure 3D, red curves). To quantify the smooth following we focus on the PFR gain for each patient measured from the “closed-loop period,” which we defined as 100-200 ms after the saccade offset. As shown in **Supplementary Fig. 1,** the PFR gain is positive and significant for both the intact (18.6% gain, CI_95_ = [18.5%, 18.6%], t_10_=4.2314, p=0.0017, BF=25.4627) and the blind fields (15.0% gain, CI_95_ = [14.9%, 15.0%], t_10_=8.4130, p<0.0001, BF>1000) but no difference between intact and blind-fields is found in this later period (t_10_=1.2041, p=0.2563, BF=0.5355). Thus, CB patients generate smooth, following eye movements for stimulus motion at the fovea after a visual response delay.

**Supplementary Fig. 1 | PFR gain in the intact and blind field of V1 stroke patients during the closed-loop period.** Plot of mean PFR gain during the closed-loop period (100-200 ms post-saccade offset) for motion targets presented in CB patients’ intact and blind fields. Individual dots represent the mean PFR gains for each participant. Error bars represent mean ± 2 SEM across subjects.

**PFR gain and respective NDR thresholds in intact and blind fields.** We performed an exponential fit to the intact visual fields to describe the dependence of PFR on the NDR threshold performance (Goodness of fit: SSE = 0.1717, R^2^ = 0.3502, and RMSE = 0.0977). As shown in **Supplementary Fig. 2,** the bold red line represents the model fit based on the intact visual field data. The dotted red lines represent the confidence interval boundaries at 95%. Out of a total of 6 blind field data points, 5 lie outside these confidence intervals, and also demonstrate respective confidence intervals on their PFR gain that are overlapping or below zero. The probability that 5 of 6 points would lie outside the 95% confidence intervals, assuming they came from the same distribution as intact fields is highly improbable (p <= 1.79x10^-6^, binomial test). We also note that the single point which fell within the range of the intact visual fields was unusual in other ways. Unlike other blind-field locations in our study, this point (Fig. 1, CB6, lower left quadrant) did exhibit significant detection performance in Humphrey visual field tests (spared Humphrey visual field performance - i.e. luminance detection), even though global motion discrimination at that location was impaired and thus selected for training (see Methods). However, we included this point to be consistent and more conservative in our analyses. Nonetheless, even when this potential outlier was included, we find quantitative evidence to support a distinction in PFR gain between intact and blind fields within the range where normal motion discrimination performance (NDR ≤ 0.35) has recovered.

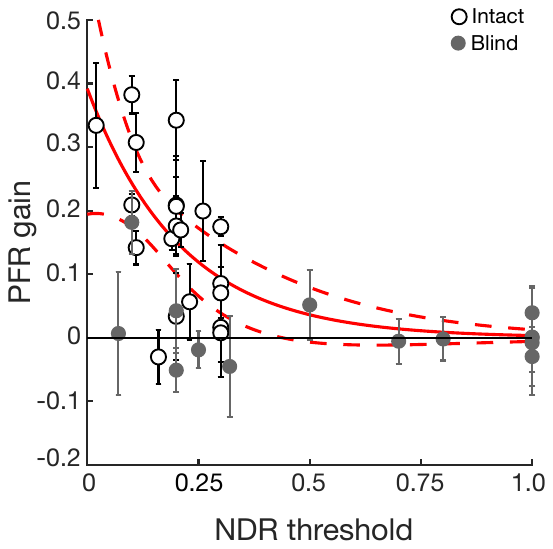

**Supplementary Fig. 2 | PFR gain as a function of NDR thresholds.** PFR gain for individual intact field locations (open circles) and blind field locations (grey circles) as a function of the normalized direction range (NDR) threshold measured at each location in stroke patients. The bold red line represents the exponential fit to the intact field, the dotted red lines represent 95% confidence interval boundaries, and the error bars on each location denote the mean ± 2 SEM.

**PFR gain versus NDR threshold pooled per subject instead by visual field locations.** As shown in **Supplementary Fig. 3**, after pooling all the fields per subject, we still observe the trend in which the lower NDR is correlated with higher PFR gain in the intact fields (R^2^=0.568, F_9_=11.8334, p=0.0074, BF=8.0855), there is no significant correlation between NDR and PFR gain in the blind fields (R^2^=0.038, F_9_=0.3552, p=0.5659, BF=2.650). Further, for NDR < 0.35, the correlation between PFR and performance remains significant for intact fields (R^2^=0.568, F_9_=11.8334, p=0.0074, BF=8.0855) but not for blind fields (R^2^=0.5406, F_2_=2.3538, p=0.2647, BF=0.7678). When we compared whether the slopes from each linear regression with respect to intact fields and blind fields under NDR ≤ 0.35, we found that slopes (-1.012 for intact and -0.1845 for blind) were significantly different from each other (t_11_=-2.6039, p=0.0245). Thus, our main findings hold when visual fields are pooled per participant to insure independence.

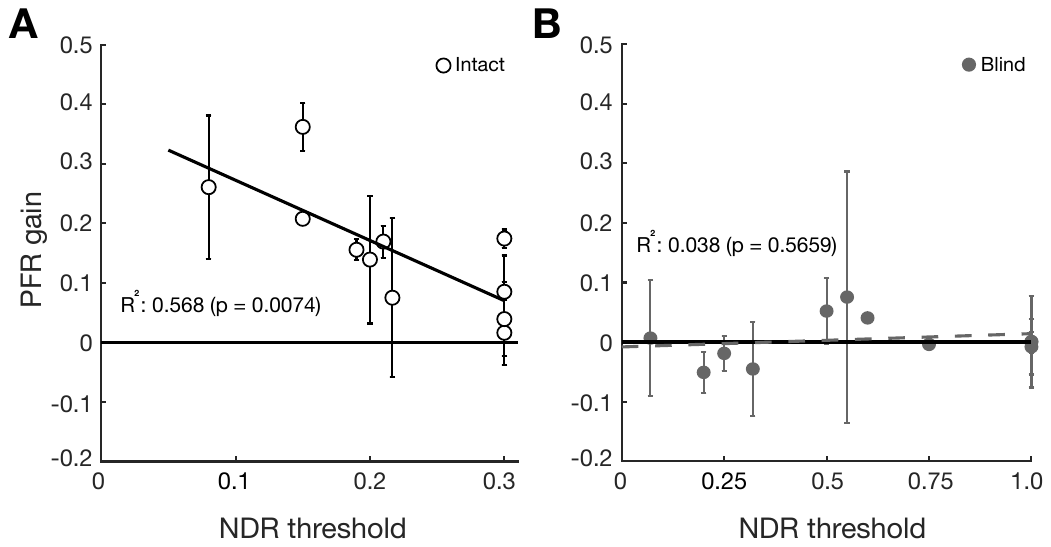

**Supplementary Fig. 3 | PFR gain as a function of NDR thresholds across V1 stroke patients.** PFR gain for individual intact field locations (Panel A, open circles) and blind field locations (Panel B, grey circles) as a function of the normalized direction range (NDR) threshold measured per each stroke patient. The error bars denote the mean ± 2 SEM.

**Luminance contrast-dependence of the PFR.** A separate group of 7 visually-intact controls were tested to determine how PFR varies with luminance contrast of the motion stimulus. Participants performed the same task as described in the main PFR experiment except that in each session, the target aperture was presented with a fixed stimulus luminance contrast between 2.5% and 50%. As shown in **Supplementary Fig. 4,** PFR gain increased steeply with stimulus contrast, starting around 5-7.5% contrast and reaching saturation quickly between 12.5-25% contrast (R^2^ = 0.9862, SSE=0.000293, RMSE=0.0086). This is comparable to neuronal responses in cortical area MT which are sensitive to low stimulus contrasts and saturate in response at roughly 10% or higher contrasts (Heuer & Britten, 2002; Kohn & Movshon, 2003; Sclar et al., 1990). Next, we considered to what extent behavior of CB patients in the blind field might reflect a response to a stimulus of effectively reduced contrast. For comparison, when CB patients performed the same test in their intact-field with 100% contrast, the observed mean PFR was 0.1448, a value that matched the performance of intact controls for contrasts ≥ 25% (**Supplementary Fig. 4**). The lowest contrast for which the PFR was significantly different from zero in controls was 7.5% (t_6_=0.3150, p=0.0202). Thus, if recovered motion perception in the blind field resembles a lower-contrast stimulus representation, then we would estimate its contrast to be less than 7.5%.

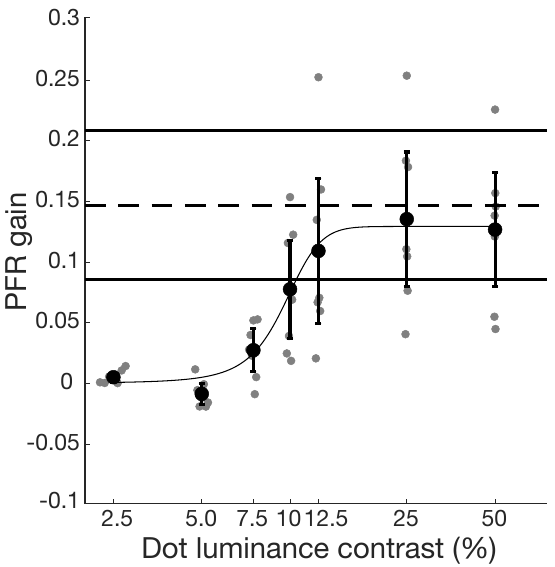

**Supplementary Fig. 4 | PFR gain as a function of dot luminance contrast in visually-intact controls.** Large black symbols and error bars denote the mean ± 2SEM of PFR gain across 7 visually-intact participants. Small grey dots represent individual participant PFR gains at each luminance level. The dashed black line indicates the mean PFR gain for 100% contrast stimuli in the intact field of our 8 CB patients, bracketed with ± 2SEM lines. A logistic function (sigmoid fit) was used to represent the best fit to the average PFR data across contrasts.

**Occipital stroke does not abolish motion-induced perceptual shifts reflected by saccade targeting.** Previous studies showed that location of an aperture is perceived as shifted along the direction of target motion contained in the aperture (De Valois & De Valois, 1991; Kwon et al., 2015; Ramachandran & Anstis, 1990). This reflects a perceptual mis-localization error along the target motion direction that also influences saccade programming by causing saccade end-points to be shifted along the target motion (Kosovicheva et al., 2014). Consistent with previous studies, our earlier study in visually-intact controls found the position of saccade end-points to be displaced along the direction of dot motion contained in a peripheral target aperture (Kwon et al., 2019). Thus, like the PFR, saccade end-points are also influenced by pre-saccadic selection of target motion, providing another measure of the predictive influence of target motion. A previous study also found that saccade end-points provide a measure that correlates well with perception (Kosovicheva et al., 2014).

Here, we asked if deviations of saccade end-points were biased by the direction of motion in both the blind and intact fields of our CB participants, and whether restoration of motion perception in the blind field influenced these saccade parameters. For each saccade, we computed the angle of the line from fixation to the saccade end-point relative to the line from fixation to the center of the target aperture. Positive angular deviations were interpreted to reflect a bias along the target motion.

As shown in **Supplementary Fig. 5**, visually-intact controls (from Kwon et al., 2019) showed a net positive saccade angular deviations that differed significantly from 0 (t_7_=11.53, p<0.001 – leftmost grey bar). This was also observed in CB patients, in intact portions of their visual fields (3.33˚, CI_95_ = [3.32˚, 3.34˚], t_10_=6.0170, p<0.0001, BF>100) - white bar in **Supplementary Fig. 5**). In fact, there were no significant differences between saccade end-point deviations between the two groups (t_17_=-0.7607 p=0.4573, BF=0.4982). In the blind-field of CB participants, saccade angular deviations were smaller than in their intact fields (t_10_=3.3586, p=0.0073, BF=7.8454), but unlike the PFRs, they were greater than 0 (1.12˚, CI_95_ = [1.11˚, 1.13˚], t_10_=3.6538, p=0.044, BF=11.7368), providing positive evidence of pre-saccadic motion
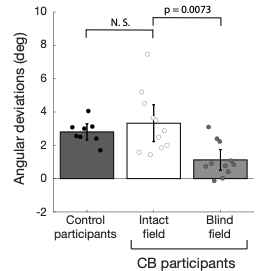
integration within the blind field.

**Supplementary Fig. 5 | Saccade angular deviations in the intact and blind field of CB participants.** Plot of mean saccade angular deviations in visually-intact controls (from Kwon et al., 2019), and for motion targets presented in CB patients’ intact and blind fields. Individual dots represent the mean saccade angular deviations for each participant. Error bars represent 2 SEM across subjects.

**Attention allocation appears intact in V1 stroke patients.**

To test for deficits in attention allocation into the blind field we ran a control experiment in two new participants. We tested if PFR might be recovered when competing distractors in the visual field were removed. This control was designed to rule out the possibility that attention impairments with respect to the blind field might emerge because attention is preferentially pulled/allocated towards stimuli that appear within intact fields, due to their higher saliency. As shown in **Supplementary Fig. 6A,** the task design was identical to the main study except for the fact that we included trials in which only a single stimulus was present in the entire visual field. A central line cue directed patients for where to saccade, and as in the main experiment, their saccades remained accurate for both intact (98.60 ± 1.15 %) and blind field (98.72 ± 0.24 %) locations. Nonetheless, as shown in **Supplementary Fig. 6B**, we did not observe a measurable PFR in the blind fields (t_3_=-1.8996, p=0.1537, BF=1.1120), while there was robust PFR in the intact fields (t_3_=3.7698, p=0.0327, BF=3.1780). PFR in intact fields was significantly larger than in their blind fields (t_3_=3.9793, p=0.0284, BF=6.8888), and the blind fields still had PFR that was not significantly different from zero (t_3=-1.8996,_ p=0.1537, BF=1.1120). Furthermore, there remained a weak correlation between PFR and NDR threshold for intact fields (R^2^=0.2818, F_10_=3.9243, p=0.0757, BF=1.0577) when compared with the blind-fields (R^2^=0.0010, F_2_=0.0019, p=0.9690, BF=0.3368) (**Supplementary Fig. 6C**).

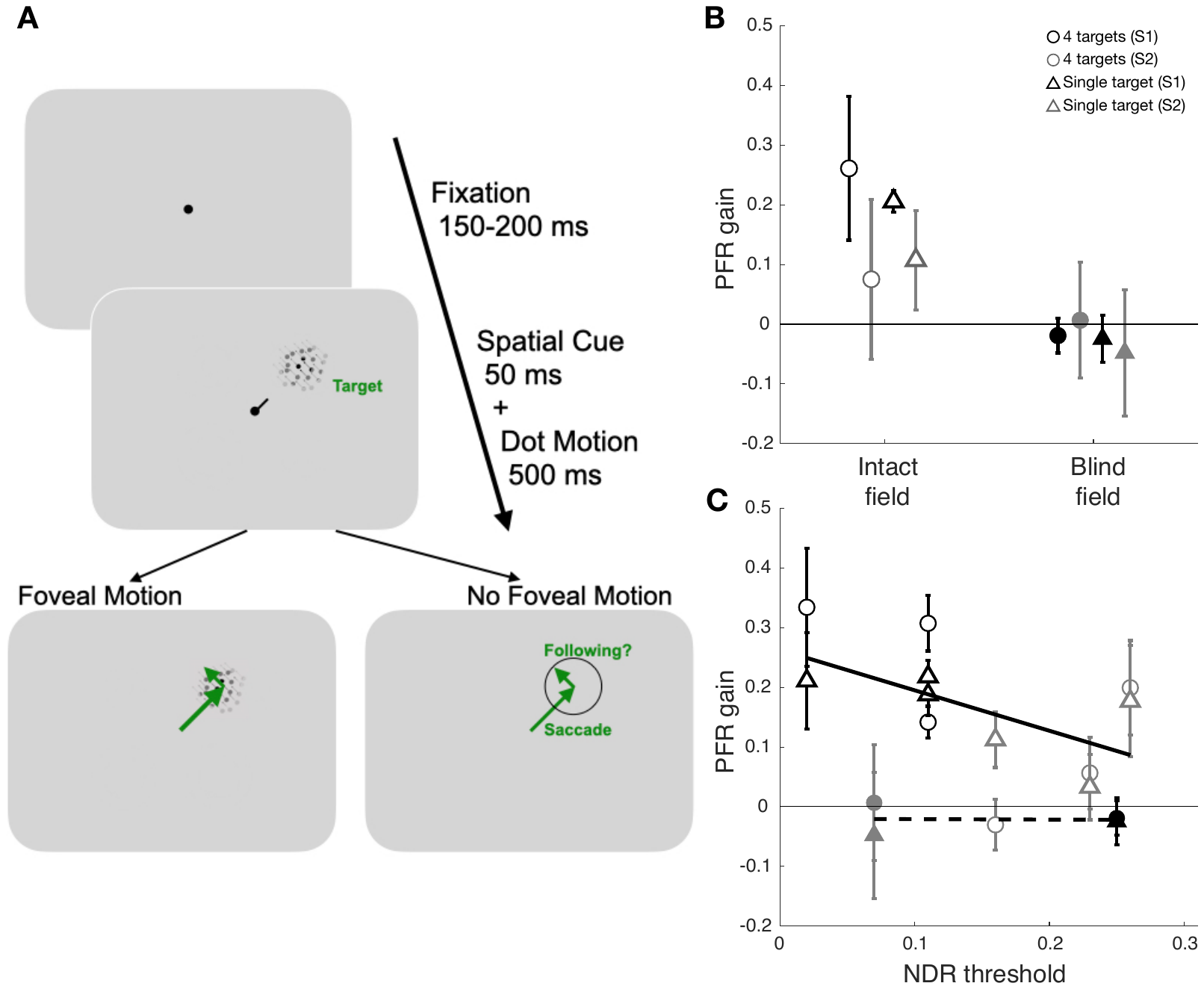

**Supplementary Fig. 6 | Experimental paradigm for measuring oculomotor functions with a single saccade target. A:** Trial sequence of assessing oculomotor behavior for a single saccade target without distractors. The paradigm followed the main experimental sequence as defined in the Fig. 2B, except only a single target that matched the spatial cue was presented at every trial. **B:** PFR gain across two patients’ intact (open symbols) and blind fields (filled symbols) in the 4 apertures experiment (circle) and single target experiment (triangle). Each color represents the respective two patients (black and grey). **C:** PFR gain as a function of NDR thresholds in each individual visual field location. The error bars denote the mean ± 2 SEM.

Thus, removing competing distractors in the other three aperture locations had little impact on the PFR (**Supplementary Fig. 6A and 6B**), suggesting the deficit was sensory in nature rather than invovling a miallocation of attention.

As a final control analysis, we also measured to what extent PFR was driven by the motion in distractor apertures. If the motion in the other three apertures produced any measurable influence on the PFR, then one might expect that influence to be stronger when saccades were made into the blind field because attention could not be properly allocated to its location, and instead would be allocated to intact field distractor locations with more salient motion targets. To test for this we repeated our PFR analyses but instead projected it onto each of the distractor aperture’s motion directions, averaging across all three of the distractor apertures. As shown in **Supplementary Fig. 7,** we did find that motion in distractor apertures contributed to a weak PFR gain when saccades were performed to a target in an intact visual field (6.85%, CI_95_ = [6.81%, 6.88%], t_10_=3.4926, p=0.0058, BF=9.4228), but for saccades made into the blind field there was no measurable PFR gain for distractor locations (1.16%, CI_95_ = [1.14%, 1.19%], t_10_=0.8594, p=0.4102, BF=0.4054), which does not support the hypothesis that greater attention was allocated to distractors for saccades made into the blind field. Althoughthe PFR gains were not significantly different when saccades were made into the intact or blind fields (t_10_=2.0156, p=0.0715, BF=1.3046). Thus, analysis of PFR for distractor apertures does not support that attention was mis-allocated for saccades made into the blind field. It seems unlikely that the lack of PFR in the blind field reflects an impairment to engage *pre-saccadic attention*.

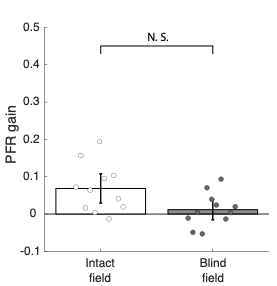

**Supplementary Fig. 7 | PFR gain in the intact and blind field of V1 stroke patients driven by motion stimuli in the distractor apertures.** Plot of mean PFR gains driven by motion in distractor apertures when CB patients made a saccade either into an intact or blind field. PFR gains driven on average by the other three distractor apertures. Error bars represent mean ± 2 SEM across subjects.
